# Cell-specific targeting by *Clostridium perfringens* β-toxin unraveled: the role of CD31 as the toxin receptor

**DOI:** 10.1101/787242

**Authors:** Julia Bruggisser, Basma Tarek, Marianne Wyder, Guillaume Witz, Gaby Enzmann, Urban Deutsch, Britta Engelhardt, Horst Posthaus

## Abstract

*Clostridium perfringens* β-toxin (CPB) is a highly active hemolysin β-pore forming toxin and the essential virulence factor for a severe, necro-hemorrhagic enteritis in animals and humans. *In vivo* and *in vitro* it exerts a remarkable cell type specificity towards endothelial cells, platelets and some leucocytic cell lines. The target cell specificity of CPB is, however, poorly understood and a receptor explaining this selective toxicity has not been identified. This has hampered further research into the pathogenesis of *C. perfringens* type C induced enteritis. Here we identify Platelet Endothelial Cell Adhesion Molecule-1 (CD31 or PECAM-1) as the specific membrane receptor for CPB on endothelial cells. CD31 expression is essential for CPB toxicity in endothelial cells and lethality in mice and sufficient to render previously resistant cells highly susceptible to the toxin. We further demonstrate, that the extracellular membrane proximal Ig6 domain of CD31 is required for the interaction with CPB and that expression of CD31 corresponds with the specificity of the toxin towards cultured cell lines. Our results thus provide an explanation for the cell type specificity of CPB and can be linked to the characteristic lesions observed a devastating enteric disease in animals and humans.

## INTRODUCTION

Pore-forming toxins (PFTs) are common virulence factors secreted by important pathogenic bacteria. Via targeting and damaging host cell plasma membranes they play important roles in the pathogenesis of many life threatening human and animal bacterial infections (Geny and Popoff, 2006; Gonzalez et al., 2008; Iacovache et al., 2008). In light of the increasing problem of antibiotic resistance, it is crucial to understand the mechanisms underlying the targeted attack exerted by these powerful bacterial weapons to develop new treatment strategies.

PFTs are typically secreted by the bacteria as water-soluble monomers that initially interact with the membrane of their target cells (Peraro and van der Goot, 2016). Surface binding leads to local concentration of the PFTs, oligomerization, and assembly of a multimeric pre-pore structure which undergoes further conformational change to insert into the lipid bilayer. Many PFTs exert cell type specific activity and this is determined by the initial binding to specific receptors on target cells. For a long time, membrane lipids were thought to function as receptors for most PFTs, however this could not sufficiently explain the cell-type and species-specificity of many of these toxins (DuMont and Torres, 2014; Peraro and van der Goot, 2016). Recently several protein receptors for important PFTs have been identified, opening new opportunities for research on the pathogenesis and therapy of bacterial infections (Rumah et al., 2015; Shrestha and McClane, 2013; Spaan et al., 2017; Tam and Torres, 2019).

*Clostridium perfringens* is a highly successful gram-positive pathogen in humans and animals. Different types of *C. perfringens* cause diseases such as food poisoning, enterotoxaemia and enteritis as well as wound infections and septicemias (Rood et al., 1997). Common to all pathogenic *C. perfringens* strains is that they affect their hosts via the secretion of highly potent exotoxins, many of which are PFTs (Petit et al., 1999; Rood et al., 2018). *C. perfringens* type C is a particularly devastating pathogen causing a severe and fatal necro-hemorrhagic enteritis (NE) in newborn animals, particularly pigs, where it can lead to high lethality rates (Shrestha et al., 2018). Human type C enteritis mainly occurred after the second world war as a disease called “Darmbrand” (Kreft et al., 2000; Zeissler and Rassfeld-Sternberg, 1949). Later on, it was identified as an important cause of childhood lethality in the highlands of Papua New Guinea, where it was finally successfully prevented after implementation of a vaccination program (Lawrence and Walker, 1976). Nowadays it is only sporadically reported in humans but remains a problem in veterinary medicine (Songer, 2010). NE is characterized by rapidly progressing hemorrhage and necrosis of the small intestine (Lawrence and Walker, 1976; Songer, 1996, 2010). *C. perfringens* type C produces α-toxin (CPA), a phospholipase, and β-toxin (CPB), a small β-pore-forming toxin of the hemolysin family (Popoff, 2014; Rood et al., 2018). Pathogenic isolates can additionally secrete many other toxins, however, CPB is the essential virulence factor of type C strains for the induction of necro-hemorrhagic intestinal lesions (Sayeed et al., 2008; Vidal et al., 2008). The toxin is secreted as a soluble 35 kDa protein (Popoff and Bouvet, 2009) and is one of the most potent clostridial toxins known (Uzal et al., 2014). It shares 28% amino-acid homology with *Staphylococcus aureus* α-hemolysin A (Hla), the prototype β-pore-forming toxin (β-PFT) of the hemolysin family (Hunter et al., 1993), which form small, usually heptameric plasma membrane pores (Berube and Bubeck Wardenburg, 2013).

In contrast to *S. aureus*, which affects a wide range of different organ systems using up to 8 currently known small pore-forming β-PFTs (Tam and Torres, 2019), *C. perfringens* type C seems to utilize mainly CPB to cause vascular damage in the intestine of susceptible hosts (Sayeed et al., 2008). We previously provided evidence that CPB primarily targets endothelial cells in the small intestinal mucosa *in vivo* (Miclard et al., 2009a; Miclard et al., 2009b; Schumacher et al., 2013). *In vitro*, human and porcine endothelial cells and thrombocytes are highly sensitive to CPB (Gurtner et al., 2010; Popescu et al., 2011; Thiel et al., 2017). Other cells with reported *in vitro* sensitivity to CPB are monocytic, lymphoblastic and lymphocytic cell lines, however the biological role of targeting similar cells *in vivo* has not been elucidated yet (Nagahama et al., 2003; Nagahama et al., 2013). Intestinal and many other epithelial as well as mesenchymal cells are insensitive to CPB (Gurtner et al., 2010; Manich et al., 2008; Nagahama et al., 2003; Popescu et al., 2011; Roos et al., 2015; Shatursky et al., 2000). Taken together these findings support the hypothesis of a cell type specific proteinaceous membrane receptor. Nagahama et al. recently suggested that the P2X purinoreceptor 7 (P2RX7) functions as a membrane receptor for CPB (Nagahama et al., 2015). The widespread expression of P2RX7 on many cell types, including most epithelia, however, cannot explain the published cell type specificity of CPB. We, therefore, hypothesized that CPB targets a membrane receptor that is expressed on endothelial cells and platelets, but absent from most other cells.

Here we identify Platelet Endothelial Cell Adhesion Molecule-1 (CD31 or PECAM-1), an adhesion molecule expressed on endothelial cells, platelets and several immune cells, as the receptor for CPB on endothelial cells. We demonstrate that CPB interaction with the extracellular membrane proximal Ig6 domain of CD31 is essential for membrane pore-formation and killing of mouse endothelial cells. Our results provide the explanation for the cell type specificity of CPB and can be linked to the characteristic lesions observed a devastating enteric disease in animals and humans.

## RESULTS

### CPB is a cell type specific toxin

To confirm our previous observation of the cell type specificity of CPB (Gurtner et al., 2010; Popescu et al., 2011; Roos et al., 2015; Thiel et al., 2017), we performed additional cell viability tests. Incubation of different human and mouse cell lines with up to 10 µg/ml CPB showed that all tested epithelial cells and fibroblasts are insensitive to CPB (Fig.1A, B, S1A, B). In contrast, all endothelial cell lines tested were highly sensitive to the toxin. The myeloid cell lines, THP-1 and U937 were also sensitive to CPB, albeit at higher concentrations compared to endothelial cells. Thus, CPB shows selectivity to endothelial and monocytic cell lines as well as platelets (Thiel et al., 2017) from different species. This lead us to hypothesize that the cell type specificity of CPB relates to a membrane receptor that is expressed on endothelial cells, platelets and leukocytes but is absent on most other cells.

**Figure 1.**
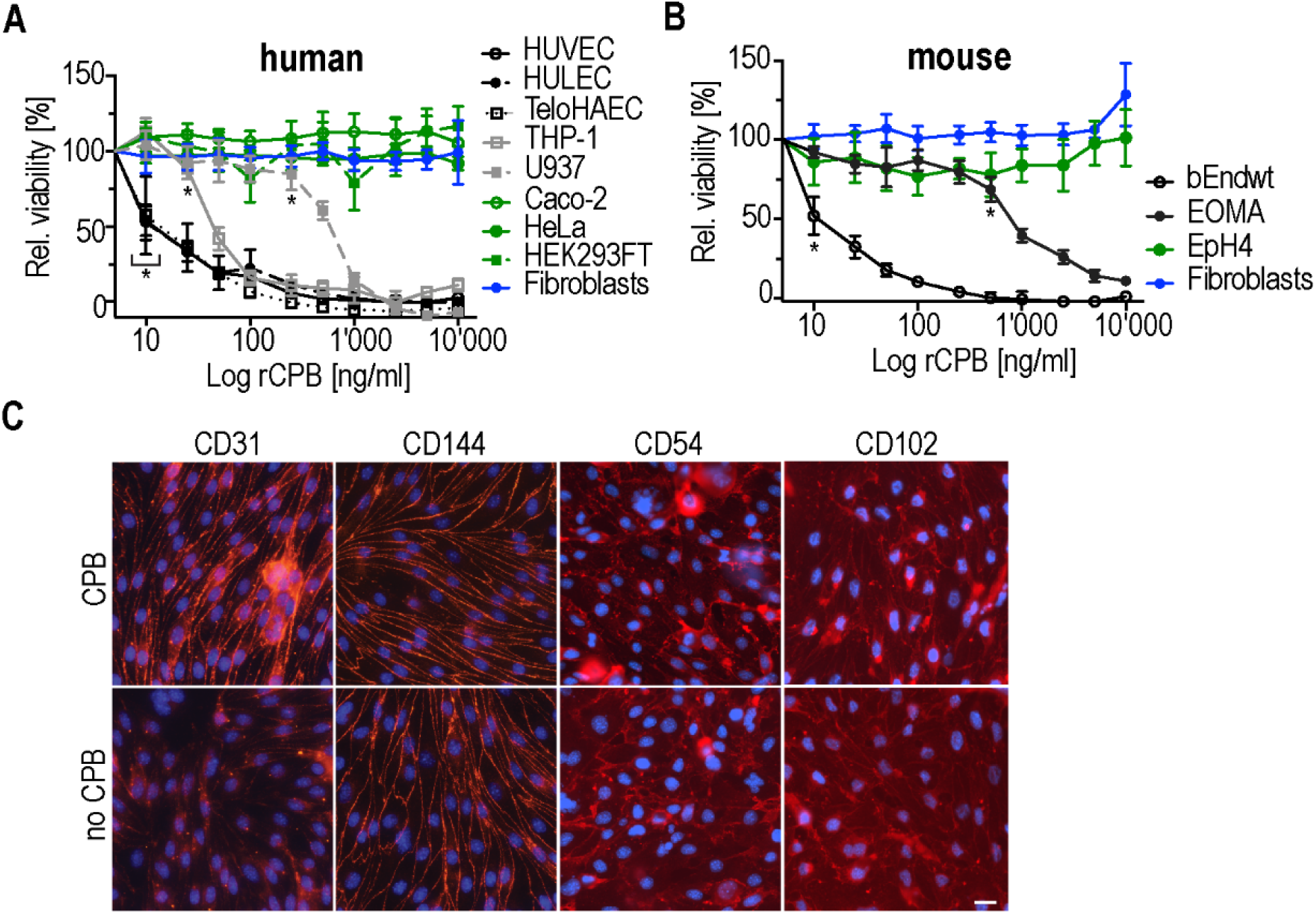
CPB is toxic to endothelial and monocytic but not epithelial cell lines. Viability of (A) human and (B) mouse cell lines in % to untreated control cells after incubation with 1 µg/ml rCPB (24 h, 37°C). N=16. Multiple comparison 2Way ANOVA, Tukey’s multiple comparison test, first values with p<0.0001 are indicated by *. (C) Immunofluorescence staining of mouse endothelioma cells (bEndwt) in the presence and absence of 1 µg/ml rCPB (10 min, 37°C) for endothelial surface molecules CD311, CD144, CD54, CD102 is shown in red. DNA stained with Hoechst (blue). Scale bar: 15 µm

### The distribution of CD31 at the plasma membrane changes after CPB exposure

We next investigated whether exposure to CPB results in a change of the distribution of endothelial adhesion proteins. Intriguingly, using immunofluorescence studies in a mouse brain endothelioma cell line (bEndwt), we found increased punctate signals for CD31 at the cell junctions shortly after CPB incubation (Fig. 1C). This effect was not observed for other endothelial cell adhesion molecules tested, such as CD144 (VE-cadherin), CD54 (ICAM-1) and CD102 (ICAM-2) (Fig. 1C).

### CPB cytotoxicity and oligomer formation depends on CD31 expression

CD31 is an important cell adhesion molecule expressed mainly on endothelial cells and on non-erythroid cells of the hematopoietic lineage which include platelets, macrophages, monocytes, neutrophils, T- and B-lymphocytes (Privratsky et al., 2010; Privratsky and Newman, 2014). Therefore, CD31 represents a plausible candidate for the cell type specific receptor of CPB. Western blot analyses showed, that all susceptible cells types tested by us express CD31 and resistant cells do not express CD31 (Fig. S1C). Importantly, also the expression levels of CD31 corresponded to the different susceptibilities of cells to CPB. To further evaluate the role of CD31 in the cytotoxicity of CPB, we performed a series of *in vitro* studies using bEndwt and CD31 knockout endothelioma cells (bEndCD31ko). In contrast to bEndwt cells, CPB neither caused cytopathic effects nor cell death in bEndCD31ko (Fig. 2A and S2). Complementary to these findings, re-constitution of bEndCD31ko cells with full length CD31-GFP completely restored susceptibility to CPB (Fig. 2A). Moreover, expression of CD31-GFP in the CPB insensitive mouse epithelial cell line EpH4 sensitized also these cells to CPB (Fig. 2B). The sensitivity of transduced cells was specific to CD31 expression, as overexpression of the related endothelial adhesion molecule CD54-GFP did not confer susceptibility to CPB (Fig. 2A, B). To determine whether CD31 influences the mechanism that leads to CPB pore formation, we monitored the conversion of CPB into SDS-resistant oligomeric forms, indicative of pore-formation. Western blotting showed that oligomer formation was only observed in CD31-proficient cells (Fig. 2C). Hence, these data show that CD31 expression is essential for the pore formation and cytotoxicity of CPB.

**Figure 2.**
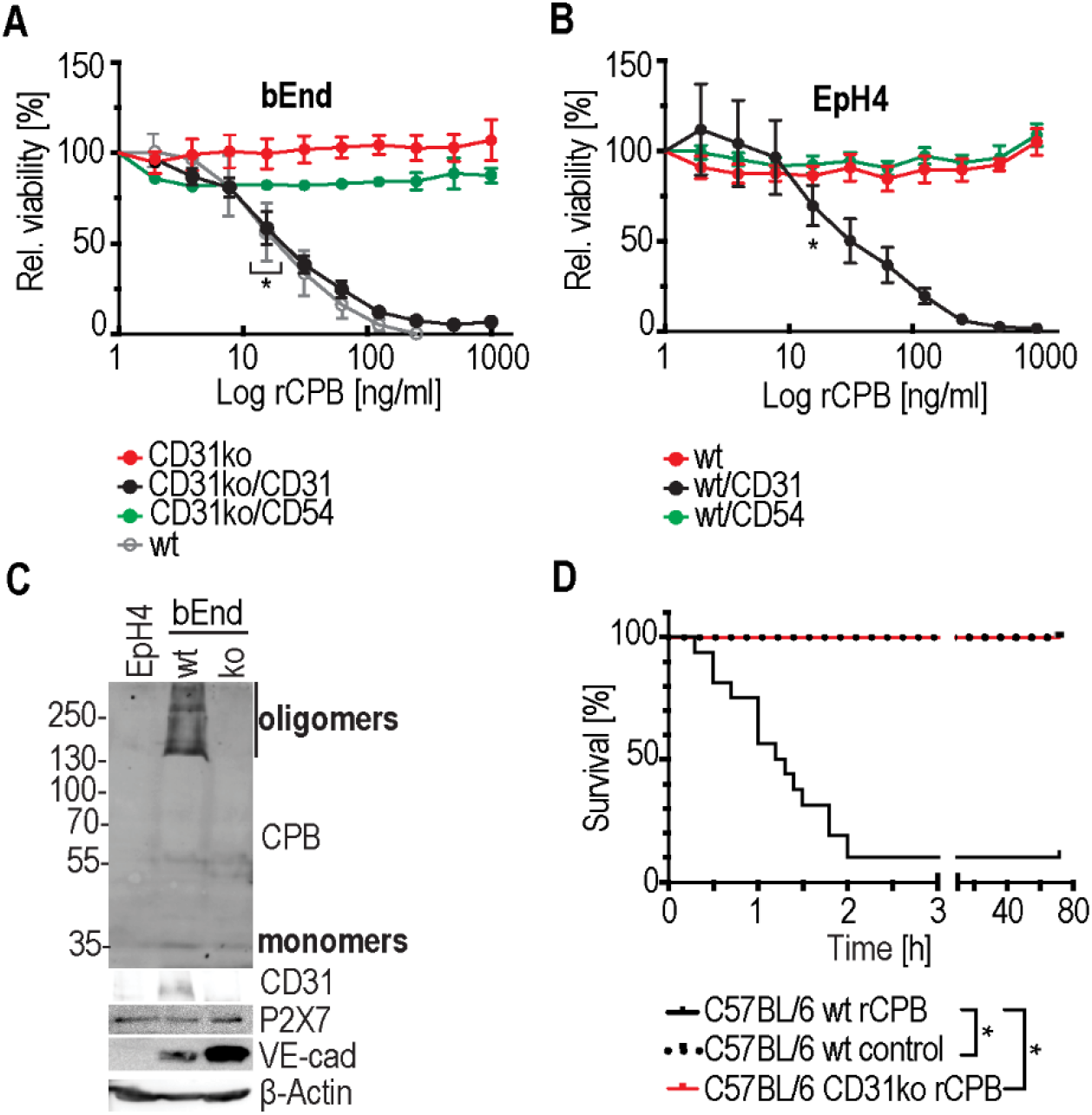
Cytotoxicity, oligomer formation, cytotoxic and lethal effects of CPB depends on CD31 expression. (A) Viability of bEndwt, bEndCD31ko, and bEndCD31ko transduced with CD31-GFP or CD54-GFP incubated with 1 µg/ml rCPB (24h, 37°C) in % to untreated control cells. N=8. Multiple comparison 2Way ANOVA, Tukey’s multiple comparison test, first values with p<0.0001 are indicated by *. (B) Viability of wild type and EpH4 cell lines stably expressing CD31-GFP or CD54-GFP incubated as in A. N=8. (C) Western blots of EpH4, bEndwt, and bEndCD31ko incubated with 8 µg/ml rCPB for 30 min at 37C° and developed with indicated antibodies. (D) Effect of CD31 depletion on the lethal activity of CPB in C57BL/6 mice. Wt and CD31 knockout mice were injected (*i.p*.) with a lethal dose (1 µg/20g). As a control, wt mice were injected with neutralized rCPB. Wild type mice (n=16), CD31 knockout mice (n=17) and control (n=13). Log-rank test, * significance (p<0.0001).

### CD31-deficient C57BL/6 mice are resistant to the lethal effect of CPB

Next, we validated the relevance of CD31 *in vivo* by investigating the effect of CD31 depletion using a mouse model of CPB toxemia. Wild type (wt) and CD31 knockout C57BL/6 mice were injected *i.p.* with a lethal dose of CPB. Wt C57BL/6 mice treated with CPB had to be euthanized within two hours after injection due to severe signs of a lethal intoxication. In contrast, CD31-deficient C57BL/6 mice were completely protected from the effect of CPB. The same was true for wt mice injected with pre-neutralized toxin as a control (Fig. 2D). These data prove that CD31 is an essential factor for CPB toxicity *in vivo*.

### CD31 directly interacts with CPB

Based on these results we investigated whether CD31 functions as a membrane receptor for CPB. Co-immunofluorescence analysis in bEndwt and bEndCD31ko cells exposed to CPB showed that CPB only binds to bEndwt cells and that CPB colocalizes with endogenous CD31 (Fig. 3A). Consistent with these findings, CPB also binds to CD31-GFP expressing EpH4 cells, whereas it does not attach to CD54-GFP expressing control cells (Fig. S3). Next, we performed *in situ* Proximity Ligation Assays (PLA) between CD31 and CPB in bEndwt and bEndCD31ko. The largest fractions of PLA interaction signals were observed at the endothelial cell junctions, consistent with our immunofluorescence analysis (Fig. 3B). In bEndCD31ko cells as well as in absence of CPB no specific signal was detected.

**Figure 3.**
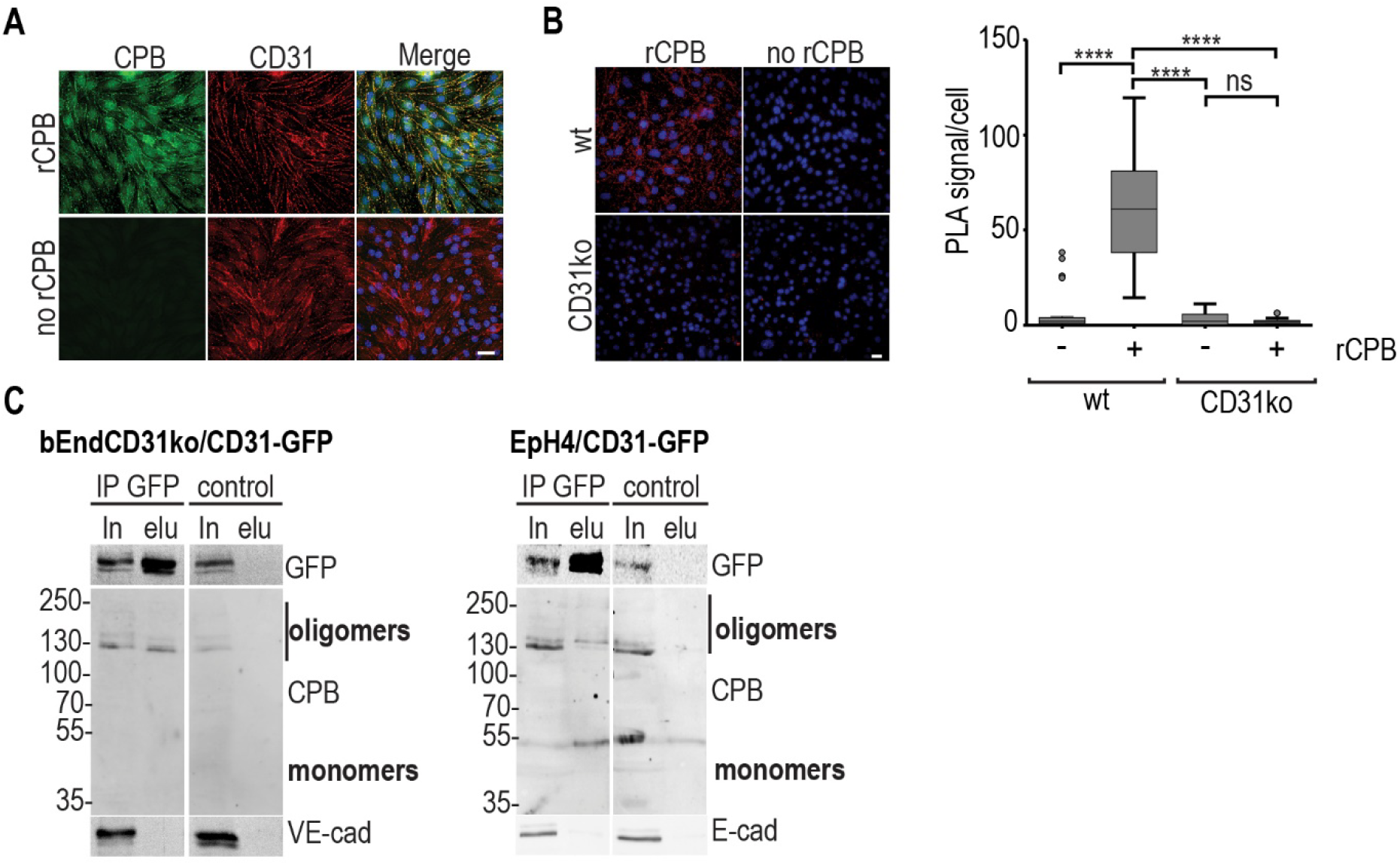
CPB targets CD31 to kill endothelial cells. (A) Immunofluorescence micrographs of bEndwt cells incubated in the presence and absence of 1 µg/ml rCPB (10 min,37°C). CPB (green), CD31 (red), DNA (blue). Scale bar: 20 µm. (B) Left panel: Maximum intensity projections of z-stacks of representative micrographs of *in situ* proximity ligation assay (PLA) of bEndwt and bEndCD31ko cells incubated in the presence and absence of 1 µg/ml rCPB (10 min,37°C). PLA signal (red), DNA (blue). Scale bar: 20 µm. Right panel: Quantification of amount of PLA puncta per cell. ANOVA test, ns (non-significant), **** (p<0.0001). N=30 (C) Immunoblots of co-immunoprecipitation experiments of CD31-GFP expressing cells incubated with 4 µg/ml rCPB and lysed (1.5% digitonin). CD31-GFP was immunoprecipitated with anti-GFP beads, anti-myc beads were used as control. Immunoblots containing 5% input (In) and 50% eluate (elu) were probed with anti-GFP and anti-CPB antibodies. E-cadherin and VE-cadherin served as a control.

To demonstrate physical interaction of the two proteins, we performed co-immunoprecipitation using anti-GFP beads to precipitate CD31-GFP protein from lysates of bEndCD31ko and EpH4 cells. CPB oligomers co-precipitated with CD31-GFP whereas no oligomers of CPB were detected in the control experiment (Fig. 3C). Reciprocal co-immunoprecipitation using anti-CPB antibodies followed by Western blotting and detection of CD31 only showed very faint signals for CD31 (data not shown), most likely because of low CD31 expression levels and consequent low levels of these complexes in the membranes of the transformed cells. We, therefore, used HEK 293FT cells, which express higher levels of CD31-GFP at their membranes. Here, we could show that the GFP-tagged CD31 co-precipitates with CPB and *vice versa* (Fig. S4).

### The extracellular domain of CD31 is sufficient for CPB cytotoxicity

CD31 consists of six extracellular Ig-like homology domains, which confer homophilic and heterophilic binding, a 19-residues transmembrane domain and a 118-residues cytoplasmic tail which is involved in signaling pathways (Privratsky and Newman, 2014). Because the initial binding of a secreted toxin to its receptor has to occur at the extracellular space, we expected that CPB interacts with the extracellular domain of CD31. To determine which domain of CD31 is involved in binding of CPB and cellular toxicity, we transduced bEndCD31ko and EpH4 cells with different GFP-tagged mutants containing the membrane bound intracellular or extracellular domain of CD31 (Fig. 4A). To exclude the effects of truncating large parts of the protein we additionally generated protein chimeras between CD31 and CD54 that belongs to the same protein superfamily of adhesion molecules. We confirmed successful plasma membrane localization of the proteins by immunofluorescence microscopy (Fig. S6). Only endothelial and epithelial cells expressing the extracellular domain of CD31 at the plasma membrane were sensitive to CPB (Fig. 4B). Expression of membrane proteins that lacked the CD31 extracellular domain but contained its intracytoplasmic domain remained resistant to CPB. (Fig. 4B). These results clearly indicate that CPB interacts with the extracellular domain of CD31 and that this interaction is sufficient to induce toxic effects in target cells.

**Figure 4.**
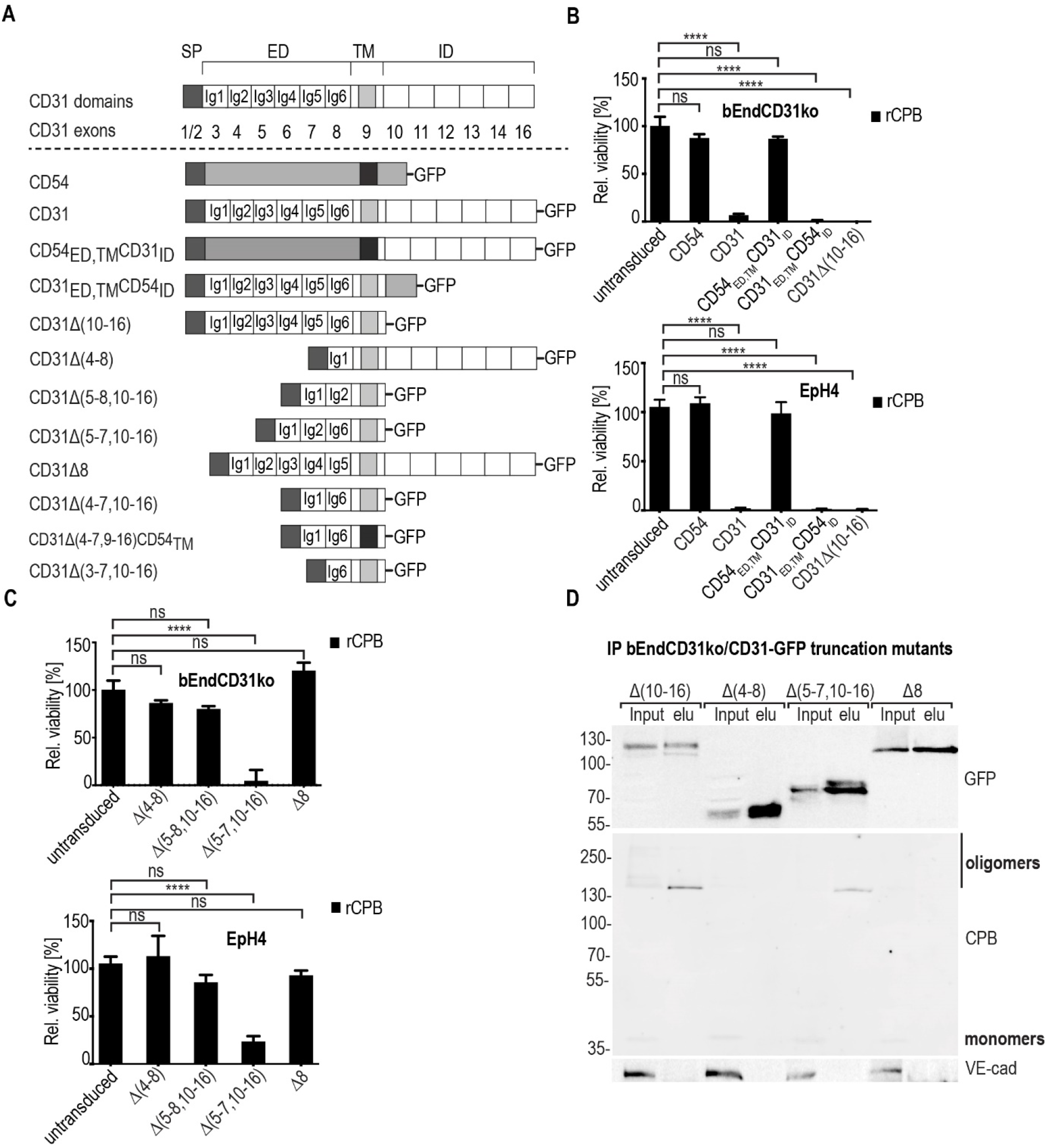
CD31 extracellular Ig6 domain is required for CPB targeting of endothelial cells. (A) Top: schematic protein domain structures of CD31 with N-terminal signal peptide (SP), the extracellular domain (ED) containing the six Ig-like domains (Ig1-Ig6), the single transmembrane helix (TM) and the intracellular domain (ID). Below, corresponding exons. Constructs below show all different GFP fusion proteins used. Deleted exons are indicated as Δ in the protein name. (B,C) Viability of cell lines expressing indicated constructs after incubation with 1 µg/ml rCPB (24h,37°C) in % to untreated control cells. 2Way ANOVA, Sidak’s multiple comparison test, ns (non-significant), **** (p<0.0001). (D) Co-immunoprecipitation experiment of bEndCD31ko cell lines expressing different CD31-GFP truncation mutants incubated with 4 µg/ml rCPB for (20 min, 37°C). IP: anti-GFP beads. Immunoblots containing 5% input (In) and 50% eluate (elu) and were probed with anti-GFP and CPB antibodies. VE-cadherin served as a control.

### CPB-cell binding and subsequent cytotoxicity requires the membrane proximal Ig6 domain of CD31

To further evaluate which Ig domain of CD31 is important for CPB-mediated cytotoxicity we generated stable cell lines expressing different CD31 extracellular truncation mutants (Fig. 4A). Plasma membrane localization of the mutant proteins was confirmed by immunofluorescence microscopy and correct protein sizes were verified by Western blotting (Fig. S5 and S6). Cell viability assays revealed that expression of the Ig6 domain containing CD31 mutants rendered cells sensitive to CPB, whereas cells expressing CD31 mutants that lack the Ig6 domain remained fully resistant (Fig. 4C). The generation of stable cell lines expressing CD31Δ(4-7,10-16) and CD31Δ(4-7,9-16)CD54_TMH_ failed, since the expression of these truncations resulted in death of transduced bEndCD31ko and EpH4 cells. We therefore transiently expressed these two constructs in HEK 293FT cells and performed additional cell viability assays. Here we could show that domains 2 to 5 are dispensable and that the CD31 membrane domain is interchangeable (Fig. S8A, B). A construct that only contains the Ig6 and the transmembrane domain of CD31 (CD31Δ(3-7,10-16)) did not localize to the plasma membrane and was therefore not used in subsequent analysis.

We next performed co-immunoprecipitation to evaluate CPB binding to different CD31 mutants. CPB oligomers only co-precipitated with CD31 constructs that contained the Ig6 domain. (Fig. 4D). Hence, Ig6 is the key domain responsible for CPB-mediated cell death.

### The conserved N-linked glycosylation site in Ig6 is not essential

The extracellular Ig6 domain of CD31 consists of 96 amino acids and is highly conserved between mouse, human and pig (Fig. 5A). It contains a conserved N-linked glycosylation site, two conserved cysteine residues that form an intra-chain disulfide bond and several conserved charged and aromatic amino acid residues. To further evaluate the specificity of the interaction between the Ig6 domain of CD31 and CPB we screened different CD31 Ig6 mutants in HEK 293FT cells. Mutation of the glycosylation site at amino acid position 540 (N540A) resulted in a mutant protein that sensitized resistant HEK 293FT cells to CPB (Fig. 5B), indicating that this glycosylation is not involved in CPB interaction with Ig6. A CD31 mutant, containing an alanine valine substitution at amino acid position 525-534 (FYKEKEDRPF 525-534 AAVAAVAAVA) did not sensitize cells to CPB (Fig. 5B). This indicates that amino acids in this region are involved in CPB binding or that the mutation affected the correct conformation of the Ig6 domain and thereby abolished the interaction between CPB and CD31.

**Figure 5.**
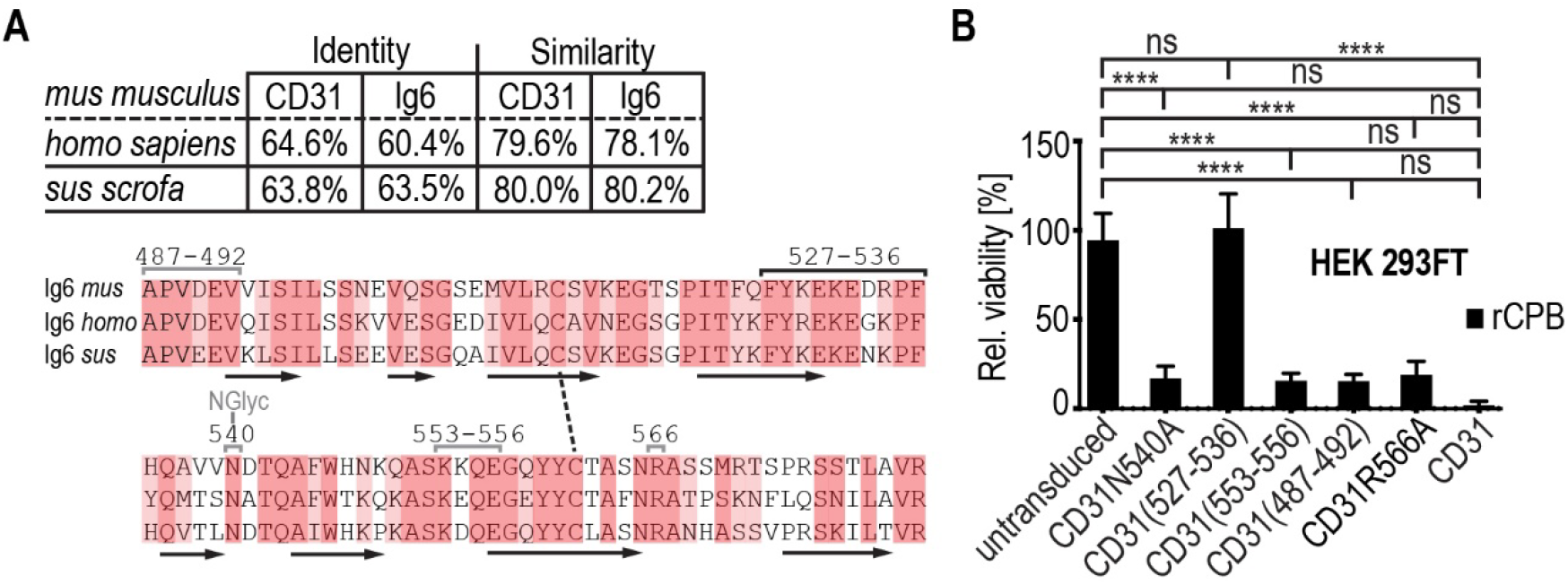
CD31 amino acids 527-536 in Ig6 are required for CPB cytotoxicity. (A) Identity and similarity values of CD31 and Ig6. Below, sequence alignments of the CD31 Ig6 domains of *mus musculus, homo sapiens*, and *sus scrofa*. Alignments were produced using EMBL-EBI server (https://www.ebi.ac.uk/Tools/msa/clustalo). Identical residues are indicated by red boxes, residues that belong to the same amino acid group by light red boxes. Mutations are indicated as grey if they were inconsequential or black if mutation diminished sensitivity to CPB. Black arrows indicate beta sheets, dotted line indicates disulfide bond. (B) Viability of transduced HEK 293FT cells expressing five CD31 Ig6 mutants (alanine and valine substitutions) after incubation with rCPB (1 µg/ml, 24h, at 37°C) in % to untreated control cells. N=12. 2Way ANOVA, Sidak’s multiple comparison test, ns (non-significant), **** (p<0.0001).

## DISCUSSION

Here we show that CD31 functions as the membrane receptor for CPB, a highly potent β-PFT that plays a major role in *C. perfringens* type C induced fatal necro-hemorrhagic enteritis in animals and humans. Previously, it has been shown that CPB is toxic to endothelial cells, platelets and several cultured monocytic, promyelocytic, lymphoblastic and B-lymphocytic cell lines (Nagahama et al., 2013; Shatursky et al., 2000; Steinthorsdottir et al., 2000; Thiel et al., 2017). Our own *in vivo* studies suggested that vascular endothelial cells in the small intestinal wall are the primary target for this toxin (Gurtner et al., 2010; Miclard et al., 2009a; Miclard et al., 2009b; Popescu et al., 2011; Schumacher et al., 2013). However, the molecular basis of this cell type specificity remained unexplained. With the discovery of CD31 as the membrane receptor, we now provide for the first time an explanation of the cell type specificity of CPB towards the above mentioned cell types.

CD31 is an important adhesion molecule highly expressed on endothelial cells, where it is a constituent of intercellular junctions (Privratsky and Newman, 2014). It is also expressed on non-erythroid cells of the hematopoietic lineage which include platelets, macrophages, monocytes, neutrophils, T- and B-lymphocytes. Due to its localization and expression on vascular endothelial cells and leukocytes that interact with the endothelium, CD31 serves a vital role in the regulation of the vascular barrier. CD31, therefore, represents a biologically feasible target for pathogens that influence local tissue perfusion to their own favor, such as *C. perfringens* type C.

Using several human and mouse cell lines we demonstrate that susceptibility of cultured cells corresponds with CD31 expression. CD31 negative epithelial and mesenchymal cells are resistant to the toxic action of CPB. Moreover, CD31 expression on endothelial cells is essential for binding of CPB to the plasma membrane, oligomer-formation, and cytotoxicity in endothelial cells. Importantly, CD31 deletion in mouse endothelioma cells and in mice abrogates CPB induced toxicity. This underlines the essential role of this membrane protein in CPB toxicity *in vivo*. Using immunoprecipitation experiments we demonstrate a physical interaction between CPB and CD31. This is supported by the co-localization of both proteins depicted by immunofluorescence and *in situ* proximity ligation. The fact that the oligomers of CPB co-precipitate with CD31 strongly suggests a direct interaction of the CPB pore with CD31 and let us speculate that they form a receptor-pore-complex. We employed different experimental approaches to determine if also CPB monomers interact directly with CD31, however, we could not enrich for monomers in elution fractions. One possible explanation could be that the rather harsh conditions that are needed to solubilize CD31 from the lipid bilayer wash off the soluble CPB monomers. Alternative explanations could be that interaction with CPB monomers has a very short half-life or that CD31 binds with higher affinity to CPB oligomers, which compete with monomers for CD31 binding.

Receptors for several β-PFTs have recently been identified, however, the exact molecular interactions between the secreted toxins and the receptor proteins still remain elusive (Rumah et al., 2015; Shrestha and McClane, 2013; Spaan et al., 2017; Tam and Torres, 2019). Here we demonstrate that CPB toxicity specifically requires the Ig6 domain of CD31. Mutational analysis of the CD31 extracellular domain revealed that membrane proximal Ig6 domain is essential for CPB action. The extracellular Ig domains 2-5, as well as the sequence of the transmembrane domain, are dispensable. Moreover, naturally resistant (CD31 negative) epithelial cells are sensitized to CPB by ectopic expression of mutant CD31 proteins containing the Ig6 domain of CD31. We were not able to investigate whether membrane bound Ig6 is sufficient for CPB binding, because mutants containing only the Ig6 domain and lacking the Ig1 domain did not localize at the plasma membrane and caused cytotoxic effects in transduced cells. The Ig1 domain promotes CD31 homophilic binding with adjacent CD31 and is essential for proper membrane microdomain localization (Newton et al., 1997). Both features might be important for CPB binding and pore formation, thus we currently cannot exclude an additional involvement of the Ig1 domain.

The Ig6 domain of CD31 has the structural requirements to facilitate direct protein-protein interaction since Ig-like domains are often recognized with high affinity (Barclay, 2003). Ig domains consist of seven to nine antiparallel β-strands arranged into two sheets linked by a disulfide bond (Alzari, 1998). To substantiate the importance of the Ig6 domain in CPB mediated toxicity, we induced alanine valine substitutions in highly conserved regions of this domain. While all mutant CD31 proteins retained normal localization to the plasma membrane, the only mutant protein that did not sensitize the cells to CPB contains mutations at position 527-536. Possibly, these residues are part of the binding epitope or the mutation altered the structure of the Ig6 domain in a way that it affected the conformation of the CPB binding site. Secondary structure prediction (JPred) on the mutated Ig6 sequence showed that the mutation did not alter position or amount of beta strands, and thus suggests that these residues are directly involved in CPB binding. The induced mutation is predicted to be located in the loop between beta strand four and five. Assuming a similar fold of Ig6 as membrane distal domains Ig1 and Ig2 (Hu et al., 2016), it would be located in a loop between beta strand four and five, a very exposed surface for potential binding partners.

Considering that a typical mammalian N-linked carbohydrate chain is almost as big as an Ig domain, glycosylation can be a major feature of these proteins and possibly involved in protein-protein interactions. Indeed, in case of aerolysin, a related β-PFT, interaction with glycan core and the N-linked sugar of the receptors are the two determinants involved in binding to the membrane (Cowell et al., 1997; Iacovache et al., 2008; MacKenzie et al., 1999). However, our data shows that the N-linked glycosylation in Ig6 is not needed for CPB cytotoxicity. Taken together these results support the notion that the Ig6 domain of CD31 contains the CPB binding epitope.

Including results from our previous studies it is clear that endothelial cells from pigs, human and mice are similarly susceptible to CPB (Gurtner et al., 2010; Popescu et al., 2011). Thus, the death of endothelial cells by CPB occurs in a species independent manner. This corresponds to the high degree of conservation of the CD31 amino acid sequence, and in particular its Ig6 domain, between these three species.

Interestingly, CD31 expression levels in different cell lines corresponded to the level of susceptibility of these cells *in vitro*. Moreover, CD31 negative cells are completely resistant to the toxic action of CPB, even at doses of up to 10µg/ml (30µM). This clearly differentiates CPB from *S. aureus* Hla, the prototype member of the hemolysin β-PFTs. Hla uses ADAM10 as a high affinity membrane receptor at low concentrations (Wilke and Bubeck Wardenburg, 2010). However, at high concentrations, Hla can induce ADAM10 independent effects on cells. At concentrations of 2 µg/ml the lipid bilayer itself seems to be sufficient for the formation of Hla heptameric pore and binding to liposomes (Bashford et al., 1996; Valeva et al., 2006; Wilke and Bubeck Wardenburg, 2010).

In conclusion, by identifying CD31 as the membrane receptor for CPB, we provide the long sought explanation for the observed host cell type specificity of CPB. Targeting of CD31 positive endothelial cells explains the early intestinal hemorrhage observed in the devastating necrotizing enteritis disease caused by CPB secreting *C. perfringens* strains. Future research should be directed into the structural basis of the toxin-receptor interaction and pore-formation to fully understand this highly specific, yet extremely potent toxin.

## EXPERIMENTAL PROCEDURES

### Reagents

Duolink® kit and PierceTM BCA protein assay kit purchased from Sigma-Aldrich. Lentiviral expression plasmid containing CD31 ORF and CD54 ORF were purchased from OriGene Technologies. EnduraTM chemically competent cells were purchased from Lucigen. Cloning kits used were Q5® Site-Directed Mutagenesis Kit and Gibson Assembly® Cloning Kit from NEB.

### Primer sequences and cloning strategy

All primers used here were ordered from Microsynth AG and sequences were verified prior to transfection by Sanger Sequencing (Microsynth AG). Cultivation of Endura^TM^ cells was done at 30°C to minimize recombination events in pLenti plasmids.

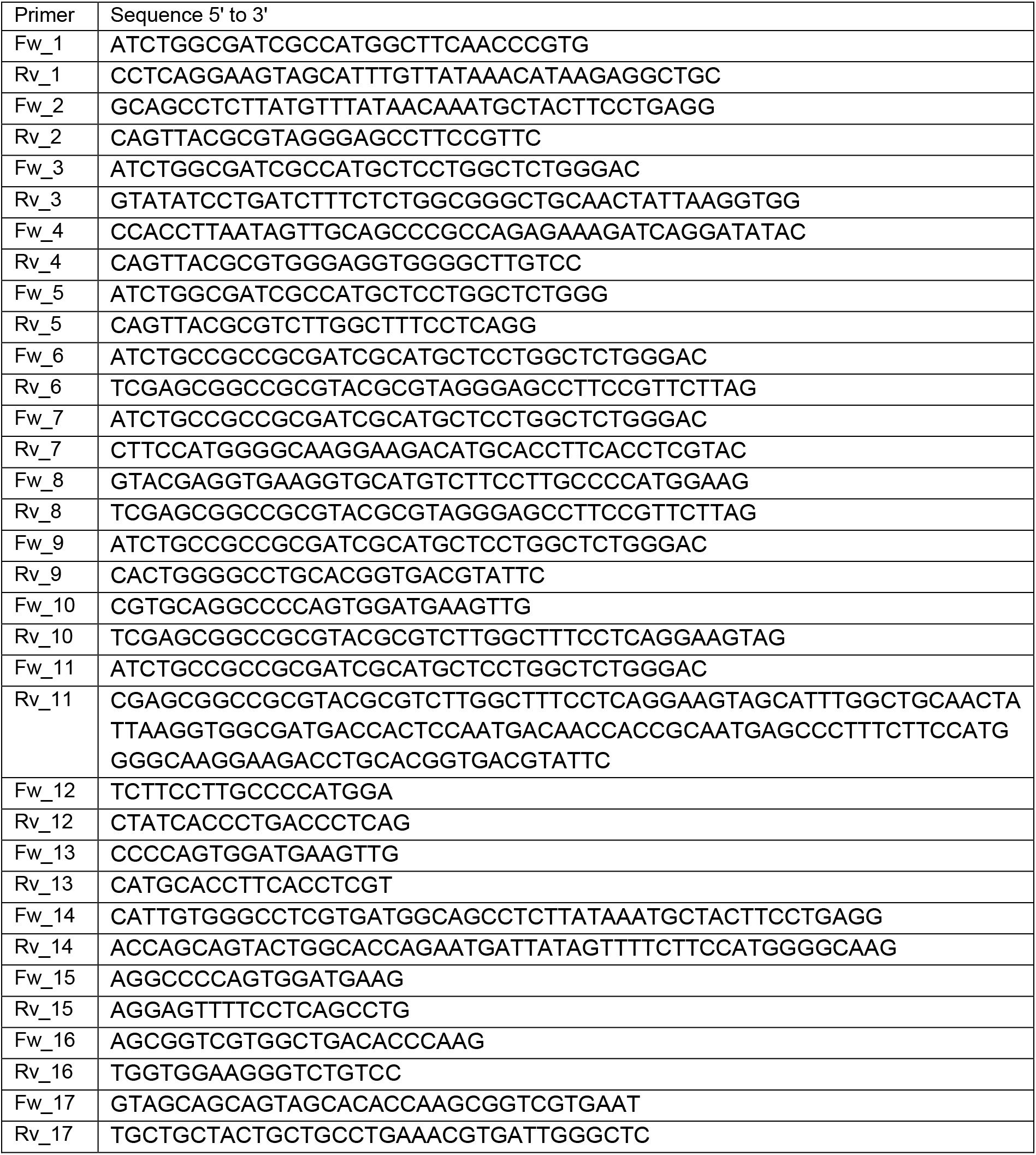
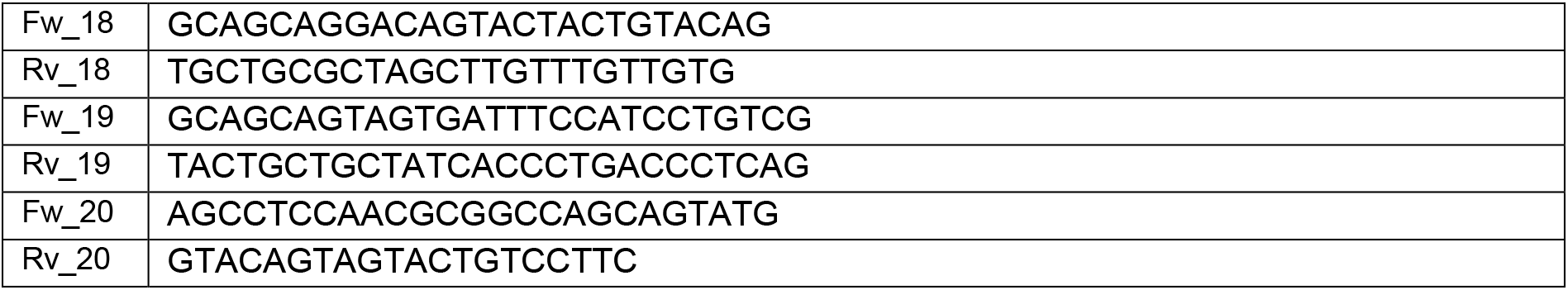

CD54_ED,TMH_CD31_ID_ was generated using pLenti plasmid that was linearized using SgfI and MluI sites. Insert containing the extracellular of CD54 part was generated from CD54 containing plasmid and primers Fw1 and Rv1, Ta of 53°C and amplification time 1min 30 sec. Insert containing the intracellular part of CD31 was generated from CD31 containing plasmid and primers Fw2: and Rv2, Ta of 52°C and amplification time 20 sec. Inserts were fused with PCR using primers Fw1 and Rv2, Ta of 56°C and amplification time 2min. The final insert was ligated into linearized backbone using T4 DNA ligase (NEB). CD31_ED,TMH_CD54_ID_ was generated using pLenti plasmid that was linearized using SgfI and MluI sites. Insert containing the extracellular part of CD31 was generated from CD31 containing plasmid and primers Fw3 and Rv3, Ta of 63°C and amplification time 1 min 30 sec. Insert containing the intracellular part of CD54 was generated from CD54 containing plasmid and primers Fw4 and Rv4, Ta of 60°C and amplification time 10 sec. Inserts were fused with PCR using primers Fw1 and Rv2, Ta of 63°C and amplification time 2 min. The final insert was ligated into linearized backbone using T4 DNA ligase (NEB). CD31Δ(10-16) was generated using CD31 pLenti full length plasmid that was linearized using SgfI and MluI sites. The insert was amplified using Q5® High-fidelity 2x Master Mix (NEB) from CD31 plasmid and primers Fw5 and Rv5, Ta of 56°C and amplification time 1 min 30 sec. The insert was ligated into linearized backbone using T4 DNA ligase (NEB). CD31Δ(4-8) was generated using Gibson cloning kit (NEB) and pLenti plasmid that was linearized using primers Fw6 and Rv6, Ta of 62°C and amplification time 4 min. Inserts were amplified using Q5® High-fidelity 2x Master Mix (NEB) from CD31 full length pLenti plasmid using primers Fw7 and Rv7, Ta1 of 63°C and amplification time of 30 sec and Fw8 and Rv8, Ta2 of 63°C and amplification time 50 sec. CD31Δ(5-7,10-16) was generated using Gibson cloning kit (NEB) and pLenti plasmid that was linearized using SgfI and MluI sites. Inserts were amplified using Q5® High-fidelity 2x Master Mix (NEB) from CD31Δ(10-16) containing plasmid using primers Fw9 and Rv9 with Ta of 63°C and amplification time of 45 sec and Fw10 and Rv10 with Ta of 60°C and amplification time 30 sec. CD31Δ(5-8,10-16) was generated using Gibson cloning kit and pLenti plasmid that was linearized using SgfI and MluI sites. Insert was amplified from CD31Δ(10-16) containing plasmid using primers Fw11 and Rv11 with Ta of 63°C and amplification time of 50 sec. CD31Δ8 was generated using Q5® site-directed mutagenesis kit, CD31 pLenti full length plasmid as template and mutagenic primers Fw12 and Rv12 with Ta of 65°C and amplification time of 3 min. CD31Δ(4-7,10-16) was generated using Q5® site-directed mutagenesis kit, CD31Δ(5-7,10-16) pLenti plasmid as template and mutagenic primers Fw13 and Rv13 with Ta of 63°C and amplification time of 4 min. CD31Δ(4-7,9-16)CD54_TMH_ was generated using Q5® site-directed mutagenesis kit, CD31Δ(4-7,10-16) pLenti plasmid as template and mutagenic primers Fw14 and Rv14 with Ta of 59 °C and amplification time of 4 min. CD31Δ(3-7,10-16) was generated using Q5® site-directed mutagenesis kit, CD31Δ(5-7,10-16) as template and mutagenic primers Fw15 and Rv15 with Ta of 65°C and elongation time of 3 min. CD31N540A was generated using Q5® site-directed mutagenesis kit, CD31 pLenti as template and mutagenic primers Fw16 and Rv16 with Ta of 63°C and elongation time of 4 min. CD31FYKEKEDRPF(525-534)AAVAAVAAVA was generated using Q5® site-directed mutagenesis kit, CD31 pLenti as template and mutagenic primers Fw17 and Rv17 with Ta of 64°C and elongation time of 4 min. CD31KKQE(553-556)AAAA was generated using Q5® site-directed mutagenesis kit, CD31 pLenti as template and mutagenic primers Fw18 and Rv18 with Ta of 60°C and elongation time of 4 min. CD31APVDEV(487-492)AAVAAV was generated using Q5® site-directed mutagenesis kit, CD31 pLenti as template and mutagenic primers Fw19 and Rv19 with Ta of 60°C and elongation time of 4 min. CD31R566A was generated using Q5® site-directed mutagenesis kit, CD31 pLenti as template and mutagenic primers Fw20 and Rv20 with Ta of 60°C and elongation time of 4 min.

### Cell culture

The endothelioma cell lines bEndwt and bEndCD31ko (bEnd.PECAM-1.2) were previously established by retroviral transduction of primary endothelial cell culture with the polyoma virus middle T-oncogene (Rothermel et al., 2005; Wong et al., 2000). HEK293FT, bEndwt, EpH4, and EOMA cell lines were cultured in DMEM medium (Gibco, product 41965-039) supplemented with 10% fetal calf serum FCS (Gibco), 10 mM Hepes pH 7.2 (Gibco), 4 mM L-Glutamine (Gibco), in the presence of penicillin-streptomycin (Gibco), grown at 37°C in an atmosphere containing 5% CO2. The media for the endothelioma cell lines bEndwt and bEndCD31ko was additionally supplemented with 1x non-essential amino acids (Gibco), 1 mM Na Pyruvate (Gibco) and 20 µM β-mercaptoethanol (Merck). HULEC 5a (ATTC® CRL 3244), Telo HAEC (ATCC® CRL 4052) and HUVEC were cultivated in Vascular Cell Basal Medium supplemented with microvascular endothelial cell growth Kit-VEGF (ATCC® PCS-110-041). Caco-2 cells, HeLa cells and NIH3T3 fibroblasts were grown in low glucose DMEM medium (Gibco, product 11054-001), 2mM L-Glutamine (Gibco), 10% FCS in the presence of penicillin-streptomycin (Gibco). THP-1 and U937 cells were grown in RPMI 1640 medium (Gibco, product 11875-093), 10% FCS in the presence of penicillin-streptomycin (Gibco).

### Production of transgenic cell lines

Transgenic cell lines were obtained by lentiviral transduction. For lentiviral production, HEK 293FT cells were transfected with third-generation lentiviral transfer vector pLenti-P2A-Puro (Origene) containing the gene of interest, a second-generation packaging vector (psPAX2), and a vesicular stomatitis virus G (VSV-G) coat envelope vector (pMD2.G) in a 5:3:2 ratio using FuGENE HD transfection reagent (Roche). The medium was exchanged 12h after transfection, and lentiviral-particle-containing medium was collected 48 h and 72 h posttransfection and filtered through a 0.4 µm filter. Transduction was performed by adding 3ml of the virus-containing medium to 40% confluent cells in a t25 flask. Cells were transduced two times, with a recovery time of 12 h between the transduction steps and selected after 36 h with puromycine (Gibco) containing media (bEndCD31ko 1.5 µg/ml, EpH4 4 µg/ml, HEK 293FT 1 µg/ml). Monoclonal cell lines were obtained by limiting dilution cloning.

### Immunofluorescence

Cells grown to confluency on 0.4% gelatin coated glass coverslips were washed with PBS and subsequently resuspended in ice-cold PBS. All further steps were performed in a wet chamber. Cells were fixed for 10 min in 4% paraformaldehyde in PBS, pH 7.2 at room temperature, and washed with ice-cold PBS. Cells were incubated in PBS containing 0.2% Triton X-100 for 10min and blocked for 45 min with 5% BSA at room temperature. Primary antibodies were gout anti-CD31 (Santa Cruz M-20, 1:100), mouse anti-CPB (Centre for Veterinary Biologics, Ames, IA 10A2, dilution 1:100), rabbit anti-VE-cadherin (Sigma-Aldrich clone Vli37, dilution 1:1’000), rat-anti CD54 (25ZC7, undiluted), rat-anti CD102 (3C4, undiluted), mouse anti-E-cadherin (Santa Cruz, DECMA-1, dilution 1:200). Alexa Fluor 596 conjugated donkey anti-goat, 488 conjugated goat anti-mouse or 594 conjugated goat anti-rabbit were used as secondary antibodies (Thermofischer, dilution 1:1’000). Cells were rinsed in ice-cold buffer and DNA was stained with Hoechst (ThermoFischer) and mounted with Fluorescent Mounting medium (Dako). Image stacks were recorded on a DeltaVision Elite High Resolution Microscope system (GE Healthcare) with Olympus IX-70 inverted microscope with a sCMOS camera, 40x and 60x Olympus Objective, and deconvoluted using SoftWorx (Applied Precision) software and images were processed with Fiji and Photoshop (Adobe). Samples in Figure 1 were analyzed on a Nikon Eclipse 80i wide-field microscope with Hamamatsu Orca R2 camera using PlanApo 40x objective (Nikon) and the Openlab 5 software (Improvision) and Photoshop were used to process the images.

### *In situ* ligation assay

Cells grown on coverslips coated with 0.4% gelatin were incubated with 1 µg/ml rCPB for 10 min at 37°C. Samples were prepared according to the immunofluorescence protocol using goat anti-CD31 and mouse anti-CPB primary antibodies and the *in situ* proximity ligation assay was performed according to the manufacturer protocol (Duolink®, Sigma-Aldrich). The coverslips were washed twice for 5 min with buffer A, followed by incubation with the PLA probes (secondary antibodies against two different species bound to two oligonucleotides: anti-mouse MINUS and anti-goat PLUS) in antibody diluent for 60 min at 37°C. After two washes of 5 min with buffer A, the ligation step was performed with ligase diluted in ligation stock for 30 min at 37°C. In the ligation step, the two oligonucleotides in the PLA probes are hybridized to the circularization oligonucleotides. The cells were washed with buffer A twice for 2 min before incubation for 100 min with amplification stock solution at 37°C. The amplification stock solution contains polymerase for the rolling circle amplification step and oligonucleotides labeled with fluorophores, which will bind to the product of the rolling circle amplification and thus allow detection. After two washes of 10 min with buffer B, the coverslips were washed with PBS and mounted with Duolink *in situ* mounting medium containing DAPI. The experiments were performed in presence and absence of CPB three times on CD31 positive cells and twice on CD31 negative cells. Duolink signal was quantified form z-stacks using a script written in Python for Fiji (Guillaume Witz. (2019, September 27). guiwitz/BernMICscripts: First release (Version v1.0). Zenodo. http://doi.org/10.5281/zenodo.3463302).

### Immunoblotting

Cells were washed with PBS and lysed with modified RIPA buffer (25mM Tris-HCl pH 7.4, 150mM NaCl, 1% NP-40, 1% Na-Deoxycholate, 0.1% SDS and complete protease inhibitor cocktail EDTA-free (Roche)) for 30 min on ice. Lysates were cleared using centrifugation (16’000 RCF, 20 min 4°C). Total protein concentration was determined using Pierce^TM^ BCA protein assay kit (ThermoFischer) and supernatants were adjusted to 2% SDS in reducing gel sample buffer and boiled for 5 min. Protein samples (40µg) were separated by 8% acrylamide Tris-glycine SDS-PAGE and transferred onto 0.45 µm nitrocellulose membranes (GE Healthcare). Membranes were blocked in blocking buffer (Odyssey® Blocking Buffer, Li-COR) and incubated overnight with the appropriate antibody at 4°C. Monoclonal antibodies used were: anti-CD31 (Santa Cruz, H-3, dilution 1:500), anti-β-actin (Sigma-Aldrich clone AC-15, dilution 1:5’000), anti-E-cadherin (Santa Cruz, DECMA-1, dilution 1:5’000), anti-CPB (dilution 1:1’000) and polyclonal primary antibodies were: rabbit anti-TagGFP (Evrogen, 1:2’000), rabbit anti-VE-cadherin (Sigma-Aldrich clone Vli37, dilution 1:1’000), rabbit anti-P2×7 (Almone labs APR-008, 1:500) and goat anti-CD31 (Santa Cruz M-20, 1:500). Detection of the antibodies was done with an Azure imaging system (c600) using IRDye secondary antibodies (Li-COR Biosciences, dilution 1:20’000).

### SDS-resistant pore formation

Cells were incubated with 8 µg/ml rCPB for 20 min at 37°C and washed twice with ice-cold PBS. Cells were lysed in modified RIPA buffer as explained above and supernatants were adjusted to 2% SDS in reducing gel sample buffer, boiled for 5 min. Protein samples (100µg) were separated by 8% acrylamide Tris-glycine SDS-PAGE and analyzed by Western Blotting.

### Immunoprecipitation

Cells were incubated with 4 µg/ml rCPB for 20 min at 37°C and washed three times with ice-cold PBS. Cells were lysed in extraction buffer (20mM Tris-HCl pH 7.4, 100mM NaCl, 25mM KCl, 5mM CaCl_2_, 10% Glycerol, complete protease inhibitor cocktail EDTA-free (Roche)) containing 1.5% (w/v) digitonin (Sigma-Aldrich) for 30 min on ice. The lysate was cleared by centrifugation (16’000 RCF, 4°C, 20 min). From the resulting supernatant an input sample (5%) was withdrawn and the rest was incubated for 45 min at 4°C in presence of anti-GFP magnetic agarose beads (Chromotek, product gtma-20), anti-cMyc magnetic agarose beads as control (Chromotek, product ytma-20) or 1h with Dynabeads^TM^ ProteinG (ThermoFischer, product 10009D) that have been preincubated with anti-CPB antibody (for 1h at room temperature). Beads were washed three times with lysis buffer containing 0.15% (w/v) digitonin and 250mM NaCl prior to elution by boiling in 2x SDS sample buffer or incubated for 20 min at 60°C (sample with Dynabeads).

### Cytotoxicity assay

Effects of CPB on cells were measured by using Resazurin assay. Cells (2 × 10^4^ cells/ml) grown to confluency in a 96 well plate were incubated with CPB Figure 1 (10 µg/ml rCPB starting concentration, 1:2 dilution steps), Figure 2 (1 µg/ml rCPB and 1 µg/ml neutralized rCPB) for 24 h. Resazurin dye (Sigma) was added to a 0.002% final concentration, incubated for 4 h at 37°C and fluorescent signal intensity was quantified using the EnSpire Multimode Plate Reader (PerkinElmer) at excitation and an emission wavelength of 540 and 612 nm respectively. For visualization of cytopathic effects, cell lines expressing indicated constructs were seeded in 8-well Nunc® Lab-Tek (Sigmaaldrich), grown to confluency and exposed to CPB (1 µg/ml) for 24h, fixed with methanol and stained with Eosin/Azur dye. Cytopathic effects were evaluated by light microscopy.

### *In vivo* toxin challenge

Animal experiments were approved by the Bernese Cantonal Veterinary Office (Animal Experiment No. BE120/16). Experiments were performed with wt and PECAM-1-/-C57BL/6 female and male mice (Duncan et al., 1999) aged 8-10 weeks and weighing 20-22 g. For analgesia, animals were injected one hour prior toxin application with 375 µg/kg buprenorphine (Temgesic) i.p. A lethal dose of 1 µg rCPB per animal (20 – 22g) in sterile NaCL injection solution was administered by i.p injection. As a control, the same dose of rCPB that was pre-incubated for 1 h at 4°C with monoclonal neutralizing anti-CPB antibody (1:5 dilution). The mice were continuously monitored and graded according to a clinical score sheet for the duration of the experiment. Animals showing severe signs of intoxication were euthanized using CO_2_ asphyxiation.

### Statistical analysis

Statistics were performed using the NCSS software (Nashville, USA, http://www.ncss.com) or GraphPad Prism 6 for Mac, (La Jolla California USA, http://www.graphpad.com).

