## Supplemental Figures for "Cell-specific targeting by *Clostridium perfringens* β-toxin unraveled: the role of CD31 as the toxin receptor"

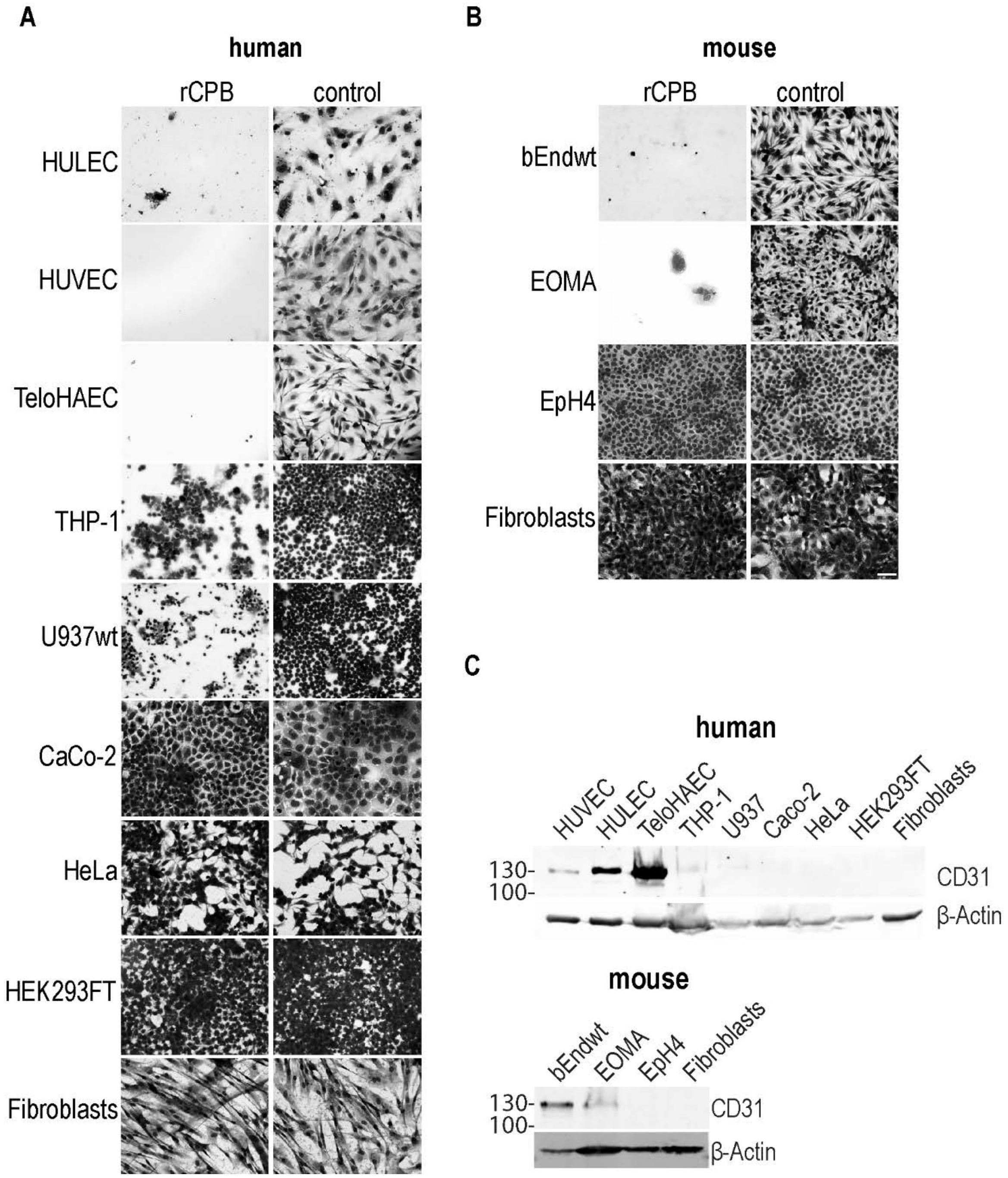


**Figure S1** Representative micrographs of (A) human and (B) mouse cell lines after incubation with 1 µg/ml rCPB (24h, 37°C). Control cells were incubated with same dose of rCPB pre-incubated with neutralizing antibody. Scale bar: 50 µm. (C) Western blot analysis of CD31 expression levels using RIPA lysates of indicated human and mouse cell lines (30µg total protein/lane). β-Actin serves as loading control.

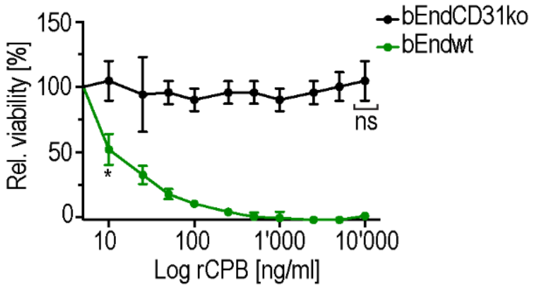


**Figure S2** Viability of bEndwt and bEndCD31ko cells incubated with rCPB at (24 h, 37°C). N=16. Multiple comparsion 2Way ANOVA, Tukey's multiple comparison test, first values with p<0.0001 are indicated by *, not significant (ns).

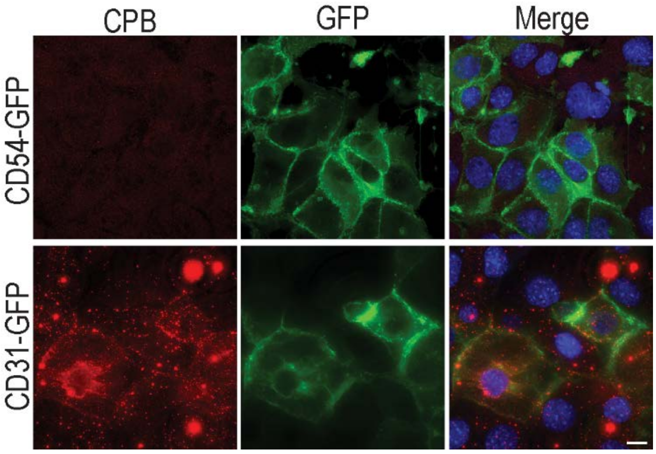


**Figure S3** Representative immunofluorscence micrographs of CD54-GFP and CD31-GFP expressing EpH4 cells incubated in presence and absence of 1 µg/ml  of rCPB (10 min, 37°C). CPB (red), GFP signal (green) and DNA (blue) in Merge. Scale bar: 10 µm.

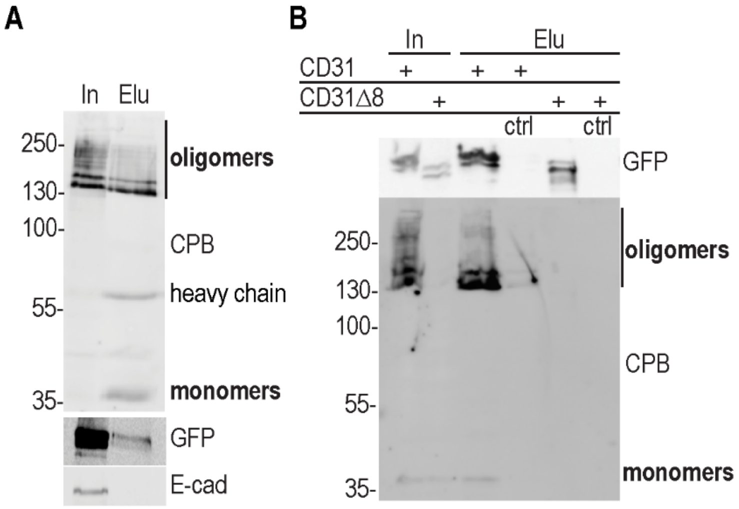


**Figure S4** (A) Co-immunoprecipitation of CPB from HEK 293FT cells stably expressing CD31-GFP. Cells were incubated with 4 µg/ml of rCPB (20 min, 37°C). CPB was immunoprecipitated from lysates using monoclonal anti-CPB antibody. Immunoblots containing 5% input (In) and 50% eluate (Elu) were probed with anti-GFP and anti-CPB antibodies. E-cadherin served as control. (B) Co-immunoprecipitation experiment of CD31-GFP or CD31Δ8-GFP from HEK 293FT lysates using anti-GFP beads in the presence of rCPB. Immunoblots containing 5% input (In) and 50% eluate (Elu) were probed with anti-GFP and anti-CPB antibodies. As controls immunoprecipitations were performed with anti-myc beads (ctrl).

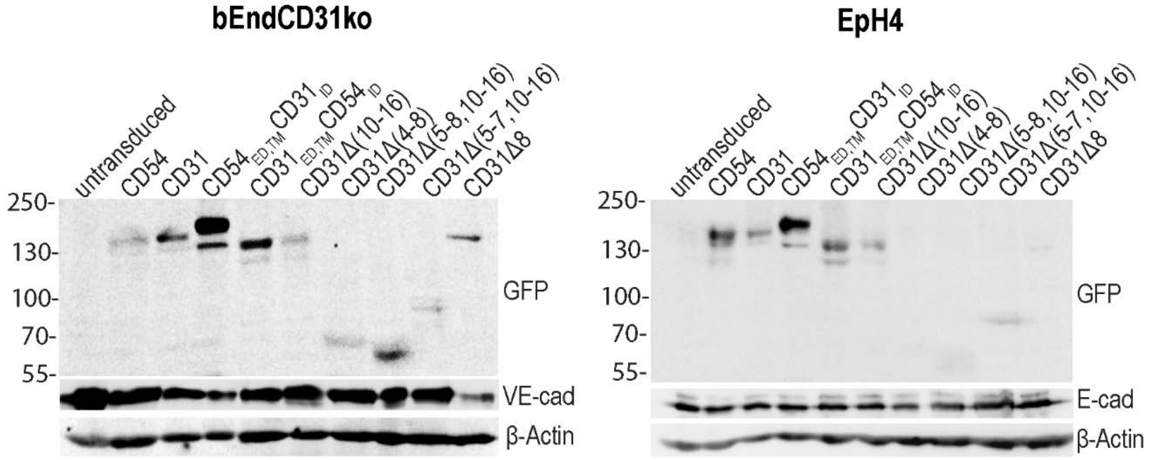


**Figure S5** Immunoblots of cell lysates from bEndCD31ko and EpH4 cell lines stably expressing indicated GFP-tagged constructs. VE-cadherin (VE-cad) and E cadherin (E-cad) included as endothelial or epithelial specific membrane markers, β-Actin as loading control.

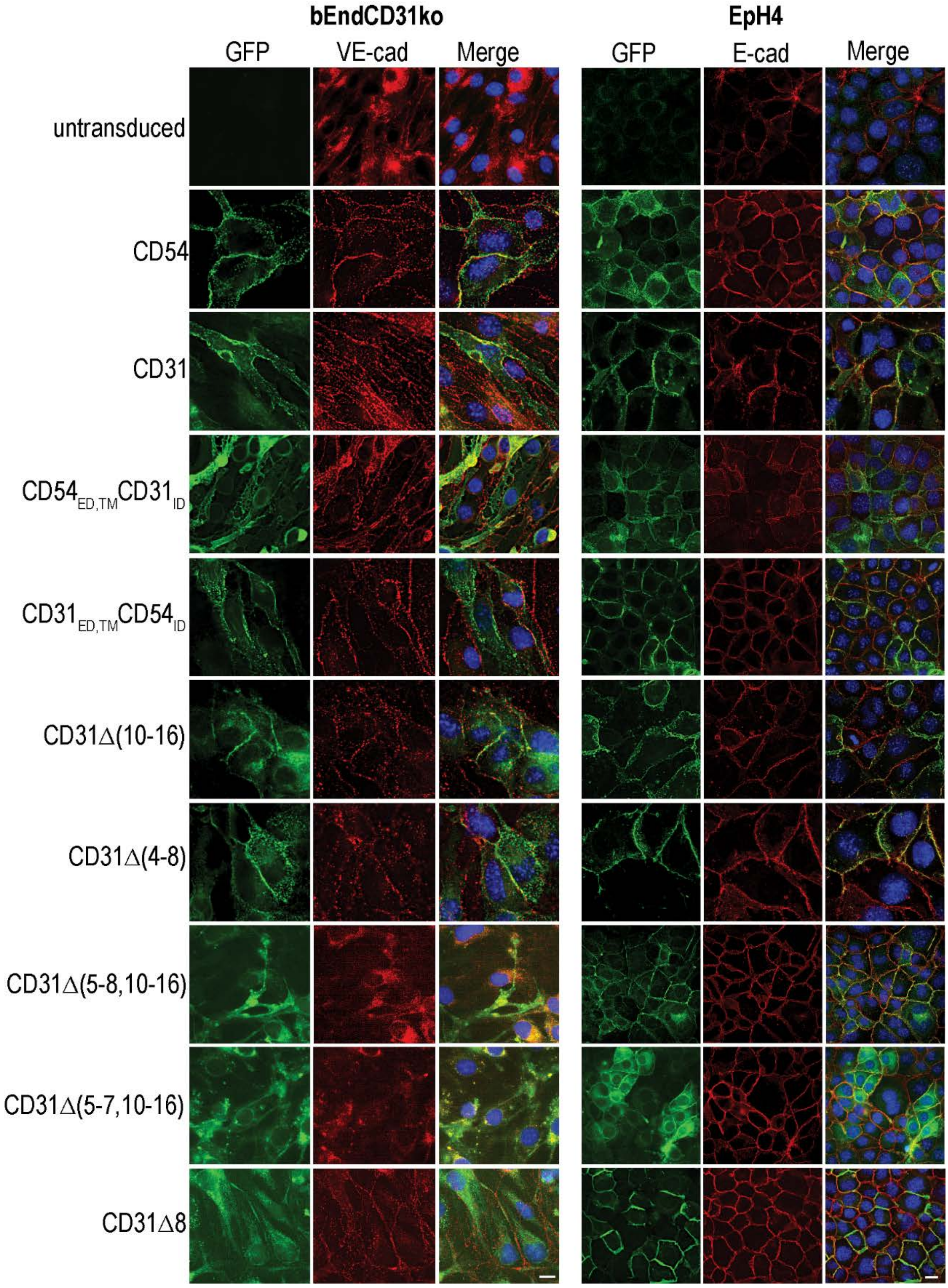


**Figure S6** Immunofluorsecence micrographs of bEndCD31ko (left panel) and EpH4 cell lines (right panel) showing plasma membrane localization of GFP-tagged proteins. VE-cadherin (VE-cad) and E cadherin (E-cad) included as endothelial or epithelial specific membrane markers. Scale bar: 10 µm.

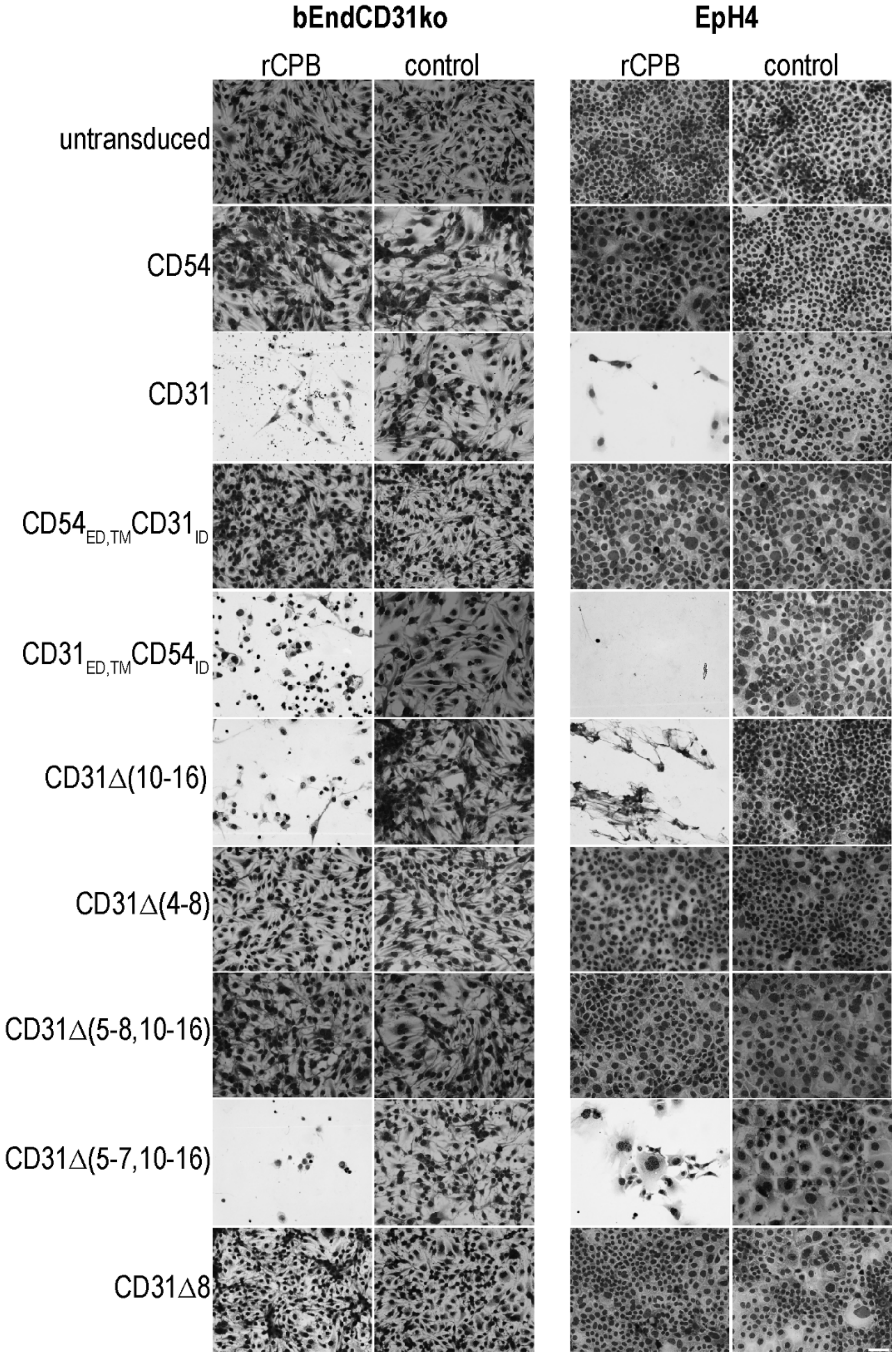


**Figure S7** Representative micrographs of cell lines expressing indicated GFP-tagged constructs showing cytopathic effect on sensitive cells after incubation with 1 µg/ml rCPB (24h, 37°C). Control cells were incubated with same dose of rCPB pre-incubated with neutralizing antibody. Scale bar: 50 µm.

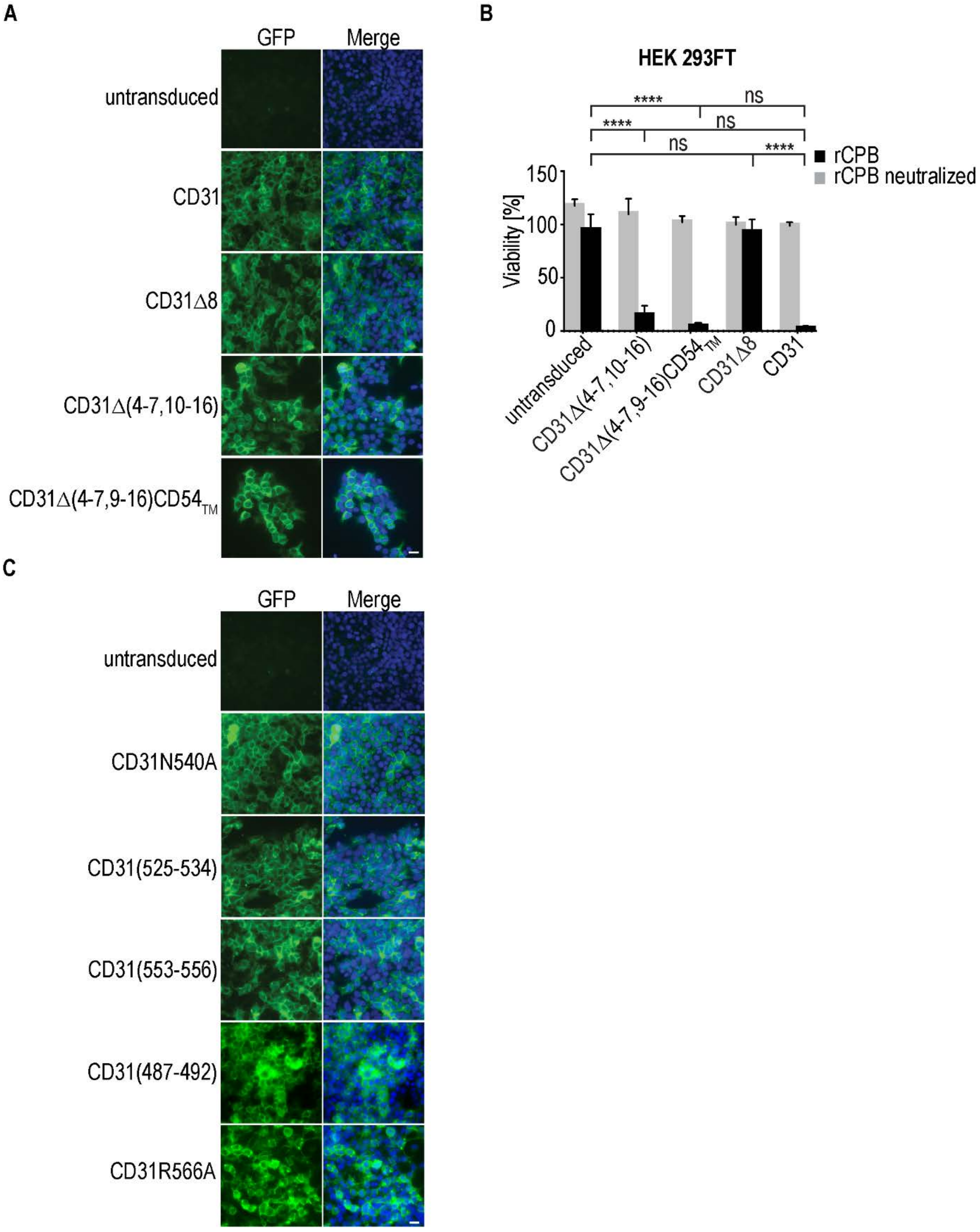


**Figure S8** (A) Fluorescence micrographs of HEK 293FT cell lines expressing indicated GFP tagged CD31 mutant proteins. Scale bar is 20 µm. (B) Relative viability of HEK 293 FT cell lines expressing indicated CD31 constructs after incbuation with 1 µg/ml rCPB (24h, 37°C) (black) or neutralized rCPB (grey) normalized to untreated control cells. N=12, 2Way ANOVA, Sidak's multiple comparison test, ns (non-significant), **** ($p<0.0001$). (C) Fluorescence micrographs of HEK 293FT cell lines expressing GFP tagged CD31 Ig6 mutants containing alanin and valine substitutions at indicated amino acid positions. GFP signal (green) and DNA (blue) in merge. Scale bar is 20 µm.
